# Conserved in 186 countries the RBD fraction of SARS CoV-2 S-protein with in-silicoT500S mutation strongly blocks ACE2 rejecting the viral spike; A Molecular-docking analysis

**DOI:** 10.1101/2021.04.25.441361

**Authors:** Amrita Banerjee, Mehak Kanwar, Dipannita Santra, Smarajit Maiti

## Abstract

SARS-CoV-2 developed global-pandemic with millions of infections/deaths. Blocker/inhibitor of ACE2 and viral-spikes Receptor-Binding-Domain RBD-blockers are helpful. Here, conserved RBD (CUTs) from 186-countries were compared with WUHAN-Hu-1 wild-type by CLUSTAL-X2 and Structural-alignment using Pymol. The RBD of ACE2-bound nCOV2 crystal-structure (2.68Å) 6VW1 was analyzed by Haddock-PatchDock. Extensive structural study/trial to introduce point/double/triple mutations in the following locations (Y489S/Y453S/T500S/T500Y)/(Y489S,Y453S/Y489S,T500S/Y489S,T500Y/Y453S,T500S/Y453S, T500Y)/ (Y489S,Y453S,T500S/Y489S,Y453S,T500Y) of CUT4 (most-effective) were tested with Swiss-Model-Expacy. Blind-docking of mutated-CUTs to ACE2 (6VW1) by Haddock-Hawkdock was performed and optimally complete-rejection of nCOV2 to ACE2 was noticed. Further, competitive-docking/binding-analyses were done by PRODIGY. Present results suggest that compared to the wild-spike, CUT4 showed extra LYS31-PHE490/GLN42-GLN498 bonding and lack of TYR41-THR500 interaction (in wild H-bond:2.639Å) with ACE2 RBD. Mutated-CUT4 strongly binds with the ACE2-RBD, promoting TYR41-T500S (H-bond: 2.0Å and 1.8Å)/T500Y (H-bond:2.6Å) interaction and complete inhibition of ACE2 RBD-nCOV2. Mutant combinations T500S,Y489S,T500S and Y489S,Y453S,T500Y mostly blocked ACE2. Conclusively, CUT4-mutant rejects whole glycosylated-nCoV2 pre-dock/post-dock/competitive-docking conditions.

## Introduction

COVID-19 spreads through the attachment viral spike-glycoprotein receptor binding domain (RBD) to the specific amino acids of Angiotensin Converting Enzyme 2 or ACE2 [1].The site specific blocking could be the major controlling point of this disease. Here, we have targeted the specific fraction of RBD, introduced one or more mutations to it and noticed that theACE2 attachment with stronger affinity. We hypothesized that, dose dependent application of our peptide might prevent the attachment of SARS CoV-2 with ACE2.In earlier and recent periods several Angiotensin Receptor Blockers (ARBs) exhibited some protecting effects. And in addition to the antihypertensive effects this drug manifested some anti-inflammatory effects also. ARBs are to some extent protective from severe acute respiratory syndrome caused by the virus [2]. Moreover, in some cases ACE-inhibitor/ARB was also found to be associated with decreased mortality [3].In contrary, intravenous infusions of ACEIs and ARBs in animal model increased expression of ACE2 receptors in the cardiopulmonary circulation. Report revealed that patients taking these therapies might be at higher risk of more viral internalization and severe disease outcomes [4]. Several studies have been conducted on the use of renin-angiotensin-aldosterone system (RAAS). The RAAS blocker and statin may have some cardiovascular benefits but whether ACE2 blockade is effective in COVID-19[5]. Our current study explains the effects of short fraction of spike protein and its mutant form in binding ACE2 to inhibit SARS CoV-2 binding. This type of work is very scanty. One study characterized bioenergetic pattern of binding-interface between the S-glycoprotein with the ACE2 receptor and tested several inhibitor peptide to SARS-CoV-2 infection [6].

The antibiotic dalbavancin was shown to bind to human ACE2 with high affinity and blocked the SARS-CoV-2 spike protein[7]. But this is not peptide-complementarily based binding and blocking which could have been more effective.

In this background, the present study was intended for the universal blocking of the ACE2 by competitive of mutant RBD fraction of SARS CoV-2 S protein. We selected the structure after the comparison of the spike proteins from 186 countries sources and then the selected amino acid T500 was mutated and found that the conserved fraction CUT4 with T500S, Y489S,T500S and Y489S,Y453S,T500Y mutations may have higher binding than the corresponding wild type. This work has great therapeutic implications.

## Materials and Method

### Structure retrieval, analysis and prediction

The X-ray crystallographic structure of ACE2 in a bonded state with nCOV2 (PDB ID:6VW1) [8]was retrieved from RCSB PDB [9]. It was with 2.68 Å resolutions and observed R-value of 0.199 (PDB DOI: 10.2210/pdb6VW1/pdb). All the amino acids involved in the binding site of nCOV2 spike glycoprotein and ACE2 was analyzed using PyMol [10]. Considering this interaction pattern as standard, other short segments were analyzed for best interaction with ACE2. The short segments were analyzed in two ways; different CUTs were prepared from tertiary structure of 6VW1 and the sequences of respective CUTs were subjected to Swiss model expasy server [11] for tertiary structure prediction.

### Docking studies

Different CUTs and their predicted structures were individually docked with ACE2. These Protein-Protein docking studies were performed using Haddock [12] and Patch dock [13] to check the binding affinity of probable structure obtained from the main and those predicted using Swiss model expasy. The results were analyzed using Pymol [10]. The docked structures were further re-docked with nCov2 to check the decreased in biding affinity of nCov2 with ACE2 RBD. Based on the complete displacement of nCOV2 spike protein from CUT pre-docked ACE2, binding affinity and smaller H-bond length of CUTs with ACE2 RBD, the best CUT was selected for further analysis.

### Mutation induction and analysis

Extensive structural study was performed to understand if introducing mutation could elevate the binding affinity of the CUT or not. Mutation sites were finalized comparing the actual interaction pattern of nCOV2 with ACE2 RBD. To strengthen the process of complete nCOV2 spike protein displacement, point mutation, double mutation and triple mutations were performed for these finalized mutation sites and there structure were predicted using Swiss Model Expasy [11].

After mutation induction, blind docking studies of different mutated CUTs with ACE2 were performed using Haddock 2.4 [12], and Hawkdock [13,14]. To obtain the best mutated structure ensuring complete nCOV2 displacement was selected through re-docking of nCOV2 with different mutated CUT pre-docked ACE2 using Haddock [12] and Hawkdock [14]. Competitive docking using Haddock was performed to analyze the competitive binding of nCOV2 and the chosen mutated structure with ACE2. Further competitive docking was performed using ACE2 (PDB ID-6VW1), the complete spike (PDB ID-6VYB) and different selected mutated CUTs. PRODIGY [15] was used to analyze the binding affinity of the competitive docking. The best mutated structures from the docking result were accepted.

### Worldwide mutation analysis

The 186 nucleotide sequences of SARS-CoV-2 genomes isolated from humans used in this analysis are available upon free registration from The Global Initiative on Sharing Avian Influenza Data (https://www.gisaid.org/) and SARS-CoV-2 spike glycoprotein gene sequence which was isolated from WUHAN –Hu-1 (COVID-19) was retrieved from the National Center of Biotechnology Information (NCBI) Biological database (https://www.ncbi.nlm.nih.gov/). Sequence comparisons with selected nucleotide sequence of SARS-CoV-2 were conducted through multiple sequence alignment using CLUSTAL X2[16]. Then spike glycoprotein gene sequences were CUT using CLC sequence viewer. These gene sequences were unique from SARS-CoV-2 spike glycoprotein gene sequence isolated from WUHAN –Hu-1. Conversion of the gene sequences into protein sequence using sequence manipulation suite tool (SMS). Sequence comparisons with unique spike protein sequences and chosen unmutated sequence were done by CLUSTAL X2 [16].The structure of the world wide spike protein sequences were predicted using Swiss model expasy [11] and the structural alignment analysis was done using PyMol[10].

### ACE1 and ACE2 Receptor Binding Domain Analysis

The nCOV2 spike glycoprotein interacts with ACE2 receptor binding domain, but not with ACE1, though two structures are highly similar and super-imposable[17]. This comparison was performed at secondary and tertiary structure level using the server of Protein Contacts Atlas (https://www.mrc-lmb.cam.ac.uk/rajini/index.html). This is a non-covalent contact based secondary and tertiary structure visualization and analysis server.

## Results and discussion

### Spike glycoprotein - ACE2 attachment site analysis

Attachment site analysis was performed using the X-ray crystallographic structure 6VW1.This PDB structure was an X-ray diffracted structure with 2.68 Å where an ACE2 interaction with spike glycoprotein was present. According to this structural arrangement, the open state ofS1 protein domain of spike glycoprotein with amino acid ALA475, ASN487, TYR489, GLN493, TYR453, TYR449, TYR505, GLY496, GLY502, THR500 and ASN501 interacted with the ACE2 surface with the amino acids SER19, GLN24, TYR83, LYS31, GLU35, HIS34, ASP38, GLU37, LYS353 and TYR41. This binding was stabilized by H-bonding ranged from 2.639-3.576Å.This interaction was facilitating the COVID-19 entry into human cell.

### Peptide screening for competitive inhibition of Spike glycoprotein - ACE2 attachment

To protect the entry of COVID-19 one strategy was taken which competitively inhibits the Spike glycoprotein - ACE2 attachment. A small segment of spike glycoprotein active site itself was administrated for competitive inhibition study. To get effective peptide sequence as well as structure, different CUTs of spike glycoprotein structures and their corresponding predicted 3D structure was analyzed for more preferable H-bonding pattern in comparison with the actual interaction as shown in Figure 1.

**Figure 1.**
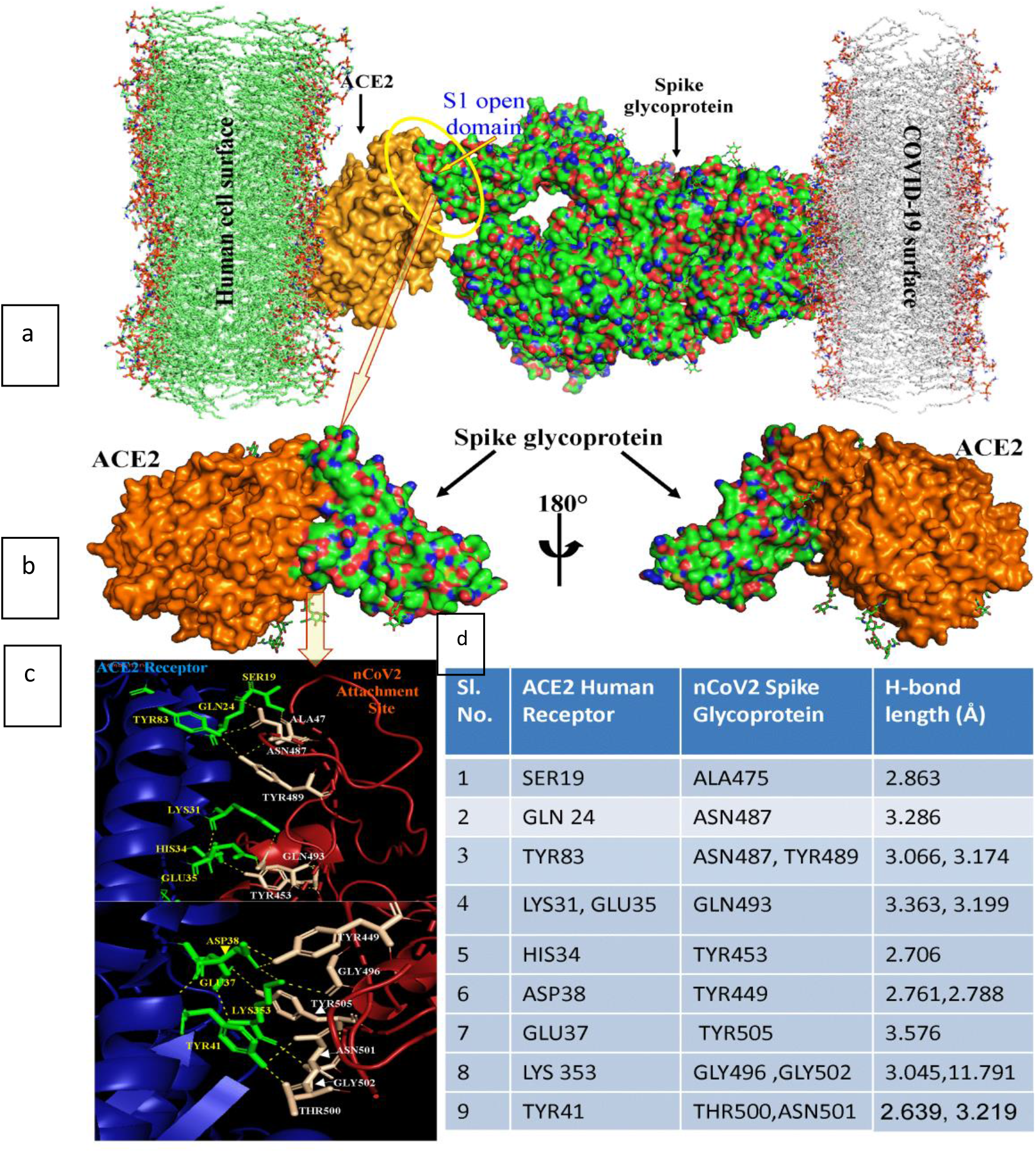
Attachment of SARS COV-2 spike glycoprotein with human ACE2 receptor. Binding interface represented in Fig a and b. Amino acid interaction pattern represented in figure c. Corresponding hydrogen bond length at each interaction was represented in figure d.

Initially at random a peptide segment of 103 amino acids was CUT from 6VW1spike glycoprotein active site location and named asCUT1 (S Fig. 1). To verify the refolding capacity of CUT1, 3D structure of selected 103 amino acids was modelled using SWISS Model (CUT1S) and compared withCUT1 structure. No such structural distortion was observed between them. Then the two segments were individually subjected to molecular docking with ACE2 from 6VW1. According to the H-bonding pattern, CUT1 showed 7 interactions with ACE2 active site ranged from 1.800- 3.416 Å (S Table 1). Whereas, CUT1S showed 6 H-bond interactions ranged from 1.7- 2.7Å.Though the affinity of CUT1 andCUT1S towards ACE2 were comparatively higher than actual (H-bond ranges from 2.639-3.576 Å), interaction HIS34-TYR453, TYR41-THR500 were absent and extra GLN42-TYR449was present in CUT1. Whereas, in CUT1S, HIS34-TYR453, ASP38-TYR449, GLU37-TYR505 and TYR41-THR500 were absent and an extra GLN42-TYR449 was present. For competitive inhibition study the CUT1-ACE2 and CUT1S-ACE2 docked structures were individually subjected to protein-protein docking with actual spike glycoprotein from 6VW1 (S Fig. 2). Partial interactive distortion of spike glycoprotein was observed except position THR500. To get complete interactive distortion other different CUTs were similarly analyzed.

**Figure 2.**
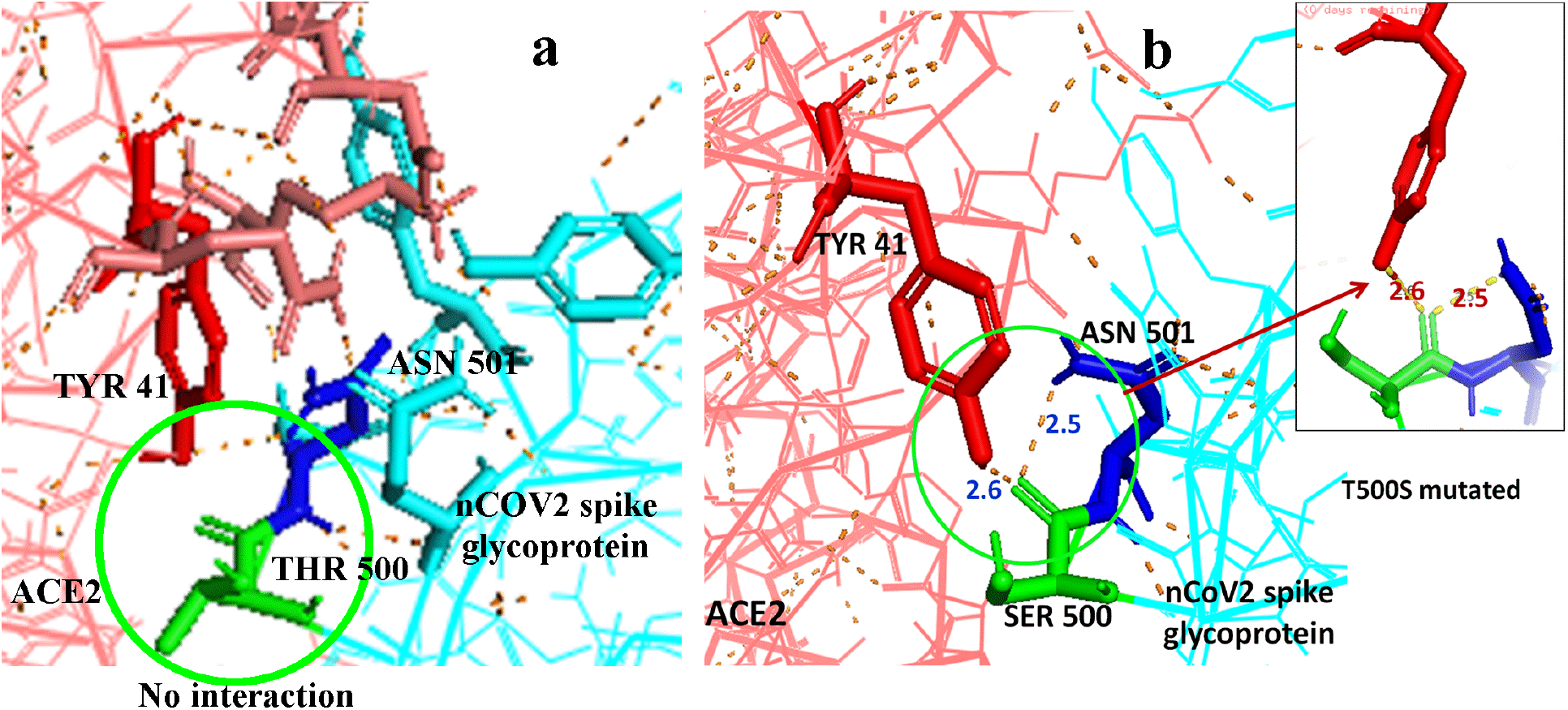
Due to H-bond based close proximity of CUT4 with ACE2 (S Table1), unmutated CUT4-THR500 did not showed and hydrogen bond with ACE2-TYR41 (a). T500S mutation only replaced the larger structure of THR with SER but mode it convenient to form H-bond with ACE2-TYR41 due to presence of same –OH group (b) and stabilized the ACE2-CUT4 mutant binding. Finally confirms the displacement of nCOV2 spike glycoprotein binding.

Structural comparison of CUT2 and CUT2S with 70 amino acids showed significant structural distortion indication loss of refolding capacity due to sequence shorting. The CUT2 did not showed HIS34-TYR453, GLU37-TYR505 and TYR41-ASN501 interactions and showed some extra bonds like GLN42-GLN48, LYS31-PHE490. But, there an imbalance was observed between CUT2 and CUT2S on interaction with ACE2 (S Table 1). No interaction was observed between CUT2S and ACE2. Multi docking of ACE2 with CUT2 and spike glycoprotein did not showed significant interaction distortion and hence it was rejected for further study.

ForCUT3, 4 amino acids were extended at the C-terminal end of CUT2 and that 74 amino acid long peptide was analyzed in a similar way. It showed some better result than CUT2. Complete structural similarity was observed between CUT3 and CUT3S (S Fig. 1). But at the interaction level, common interaction of SER19-ALA475 was absent. Like others HIS34-TYR453 and GLU37-TYR505 were also absent in both CUTs. Whereas, the important interaction TYR41-ASN501, THR500 was absent in CUT3S (S Table 1). Better result was observed in multi docking among CUT3-spike-ACE2 and CUT3S-spike-ACE2. Complete distortion of spike was observed while using CUT3. For CUT3S, spike interaction at the nearby location was observed except the active site (S Fig. 2). Though CUT3 showed better docking result, further peptide screening was performed as CUT3 devoid of proper active site interaction pattern (S Table 1).

CUT3 was extended at both C & N terminal end and CUT4 with 84 amino acids was selected for further study. CUT4S also showed proper refolding and structural similarities with CUT4 (S Fig. 1). According to the CUT4 and CUT4S docking analysis with ACE2 revealed 9 and 11 interaction sites, among them 7 and 8 were common for CUT4 and CUT4S respectively in comparison with spike glycoprotein.CUT4 showed extra LYS31-PHE490 and GLN42-GLN498 interaction. In addition to that, CUT4S showed TYR41-ASN501 and GLN325-ARG439 interaction. Multi docking with CUT4-ACE2-spike showed interaction withspike protein whereas CUT4S-ACE2-spike doesn’t show any interaction with the spike glycoprotein at the specific site of attachment in comparison toCUT4.The interaction of TYR 41 -ASN501 and GLN 325-ARG439 in CUT4S-ACE2 (S Table 1) can be held responsible for blocking the site for the attachment of spike glycoprotein as compared to CUT4(S Figure1). According to different scores analysis during molecular docking with HADDOCK, best HADDOCK score of −121.9+/−5.6 was observed for CUT4S. Which also showed Van der Waals energy and Electrostatic energy value of −74.2 +/− 1.0 and −167.6 +/− 4.4 (S Table2).

ThereforeCUT4S was taken for mutation study and the site of interaction in CUT4S-ACE2 preventing the interaction with spike glycoprotein was taken as the targeted site for mutation induction.

### Mutation induction and competitive inhibition of Spike glycoprotein - ACE2 attachment

To further enhance the inhibition of spike glycoprotein and improve the binding of CUT 4 with ACE2 receptor, mutation was induced at the positions ofTYR489, TYR453and THR500.These three amino acids of main nCOV2 spike glycoprotein showed interactions withTYR83, HIS34 and TYR41 at the ACE2 surface according to 6VW1 (Fig. 1).Whereas, in different CUT analysis, abovementioned interactions were not found(S Table 1). Threonine (THR) possess a hydroxyl (–OH) group at its R group. At location THR500, that –OH group forms hydrogen bond with the R group of TYR41 of ACE2. As Serine (SER)and Tyrosine (TYR) also have R specific hydroxyl (–OH) group, substitution with SER and TYR were made at the selected locations. As according to the previous study, the overall hydrogen bond length of CUT4 with ACE2 was less than the actual spike interaction (S Table 1, Fig. 1), SER was introduced at the positions of 489 and 453 to avoid the steric hindrance due to large R group of TYR. And both SER and TYR were separately introduced at the location of THR500. Different permutation and combination of single (4 structures), double (5 structures) and triple (2 structures) position mutations were analyzed (S Fig. 3).In total 11 structures were predicted using Swiss Model and individually docked with ACE2. In single mutationY489S, Y453S and T500Y did not show proper interactions (S Table 3)and hence were rejected for further study. Whereas, T500S mutation showed hydrogen bonding with TYR41 of ACE2, which mimicked the actual ACE2-nCOV2 spike-binding features (Fig. 2).This interaction was initially lost in case of CUT 4 and hence SM3 was considered for further study. Double Mutation 1(DM1) with Y489S Y453S showed no interaction at the mutated site and comparatively lesser interacting sites as compared to SM3 hence were rejected for further study. DM2 with Y489S T500S showed same number of interacting sites as compared to SM3. Here mutated SER 500 bonded with TYR41 with a bond length of 1.8Å hence was also considered for further studies. DM3 with Y489S T500Y, DM4 with Y489S T500Y and DM5 with Y453S T500Y all showed fewer interacting sites as compared to DM2 (S Table 3) with no interaction at the mutated sites (S Fig. 2) and hence were all rejected for further studies. Triple Mutation 1(TM1) with Y489S Y453S T500S mutation also showed no interaction at the mutation site hence not considered for further study. TM2 with Y489S, Y453S, T500Y though had same number of interacting sites as TM1(S. Table3) showed interaction of TYR500 with that of TYR41 with Hydrogen bond length 2.6 Å(S Fig.2). Hence the selected three sets were considered for further studies. Finally, three combination of mutation SM3, DM2 and TM2 individually showed mimicked interaction with ACE2 in competition with nCOV2 spike protein.

**Figure 3.**
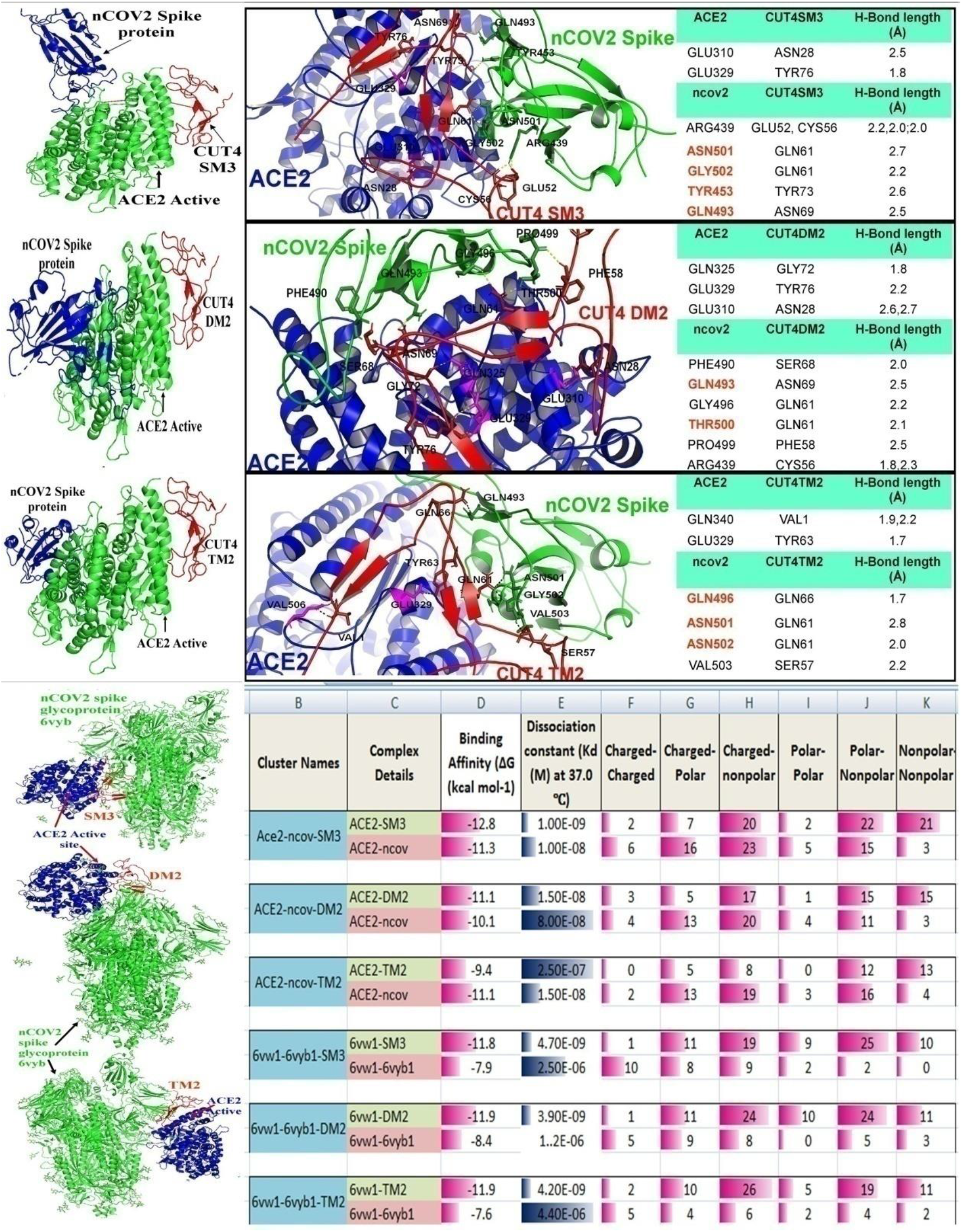
Competitive interaction of nCOV2 spike glycoprotein and Different selected mutatedCUTs with ACE2. The selected mutated CUT4s showed highest affinity to ACE2 and that attachment completely distorted the nCOV2 spike glycoprotein from ACE2 attachment (a, c, e and g). According to attachment site analysis, mutated CUT4 interacted with different active site molecules on nCOV2 spike glycoprotein, indicating their distortion with ACE2 (b, d and f). Figure h indicating the binding affinity of ACE2 with nOCV2 individually in presence of SM3, DM2 and TM2. Where, SM3 and DM2 showed more affinity to ACE2. Whereas for complete spike glycoprotein (6vyb1), Mutated CUT4 get more preference in ACE2 attachment as they showed binding affinity of −11.8, −11.9 and −11.9 respectively. And all SM3, DM2 and TM2 showed less dissociation from ACE2 in comparison to nCOV2 spike.

The docked structure ACE2-CUT4 SM3, ACE2-CUT4 DM2 and ACE2-CUT4 TM2 were further used for multiple docking with nCOV2 spike protein (Fig. 3). Before mutation an uncertain inhibition of nCOV2 spike protein interaction with ACE2-CUT4 was observed. Where nCOV2 spike protein showed partial interaction with ACE2 active site in presence of CUT4 main, but completely detached from ACE2 active site in presence of CUT4S. This uncertainty was further channelized towards certainty through specific mutations at above mentioned position. For all the selected mutations SM3, DM2 and TM2, nCOV2 spike protein showed complete displacement from ACE2 active site. Which indicated that, pre-administration of these selected mutated peptides in physiological condition, will inhibit the nCOV2 spike protein interaction with ACE2 (Fig. 3 a, c and e).

To understand the competitive interaction of nCOV2 spike protein and individual mutated peptideCUT4 SM3, CUT4 DM2 and CUT4 TM2 with ACE2, a multi ligand docking study was performed with ACE2 using HADDOCK. Results represented that, ACE2 is partially docked by both of nCOV2 spike and CUT4 mutants at the active site. In addition to that, some of the nCOV2 spike proteins active site amino acid residues were blocked by CUT4 mutants (Figure 3b). For single mutation, ASN501, GLY502, TYR453 and GLN493 of nCOV2 were blocked by CUT4SM3. These amino acids stabilize the interaction with ACE2 at the middle and lower position. SimilarlyCUT4DM2 and CUT4TM2 were also showed significant interactions with nCOV2 spike active site (Fig. 3d and f). This competitive interaction results that if sufficient quantities of CUT4 mutants are available at the physiological system, they will destabilize the ACE2-nCOV2 spike interaction. For further confirmation, a complete glycosylated nCOV2 spike protein was selected for ACE2 interaction study in presence of CUT4 SM3, CUT4 DM2 and CUT4 TM2 individually. Results indicated the complete ACE2-nCOV2 inhibition in all cases as shown in figure 3g.The competitive interaction study was further analyzed through Binding affinity (Figure 3h). Figure h indicating the binding affinity of ACE2 with nOCV2 individually in presence of SM3, DM2 and TM2. Where, SM3 and DM2 showed binding affinity of −12.8 and −11.1 in comparison with ACE2-nCOV2 interaction with the binding affinity value of −11.3 and −10.1 respectively. On the other hand the complete spike glycoprotein binding affinity was low or moderate comparative to ACE2-CUT4 mutant binding. This result indicated that different CUT4 mutant can able to inhibit ACE2-nCOV2 spike protein interaction.

Finally, the presence of CUT4 was studied among 186 COVID-19 spike proteins from 105 different countries. Those sequences were subjected to common sequence shorting. The result indicated 24 unique sequences among all (Fig. 4a). According to alignment, one sequence from Finland showed a substitution of S instead of A. But this change has not reflected in T500 position upon which the complete displacement of nCOV2 spike is dependent. Other common conserved pattern from different countries like; ESTONIA, Latvia, Hong Kong, Costarica, Iran, Mexico, Mongolia, Japan, Italy, Egypt, Ireland, Denmark, Germany, France, India, DRC, Serbia, Pakistan, England and Wuhan (wild type)also depicted the conserved T500 (Fig. 4b and c). It is indicating the universality of our CUT4 mutants.

**Figure 4.**
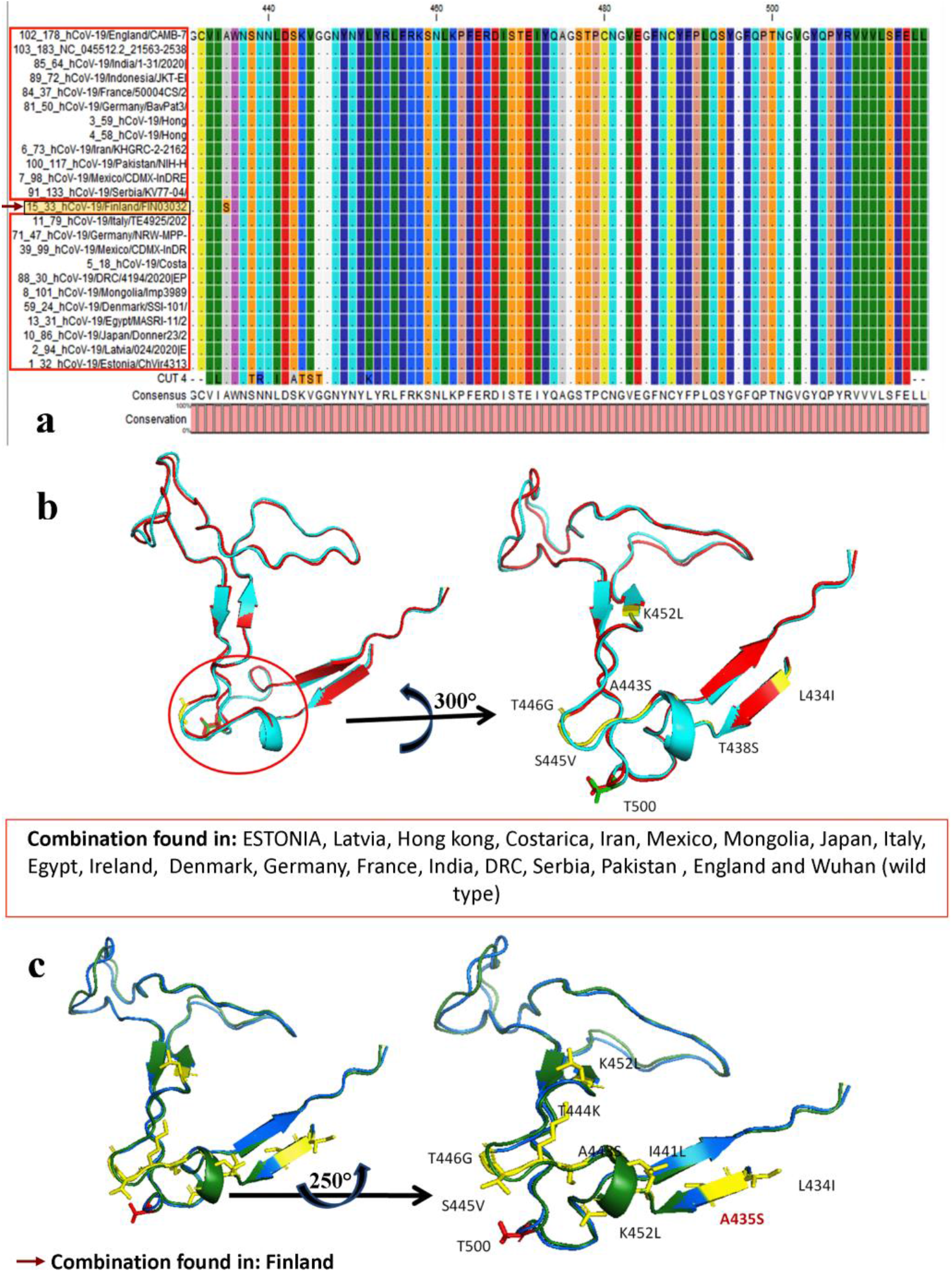
CUT4 sequence alignment with 24 unique spike proteins from different countries. These 24 unique sequences were selected from 186 spike protein sequences representing 105 countries (a). Sequence diversity was observed in sequence FIN30302 where one Ser was observed at alignment position 435 and in CUT4 within alignment position 452. Whereas, all other sequences showed identical sequences over the CUT regions. The sequence diversity with CUT4 did not reflected at the T500 position for both figure 4b and 4c. Which indicated that mutated CUT4 can able to compete with all type of COVID −19 spike glycol proteins throughout 105 countries in the world.

According to literature, ACE2 is the sole receptor of COVID19 spike glycoprotein but not ACE1, though two structures are similar. In reference to that we have studied the RBD of ACE1 and ACE2 (Fig. 5). Two structures were similar at sequence, secondary and tertiary level also. Only, the Helix1 (H1) of ACE1 started from residue 14 but H1 of ACE2, started from residue 41.ACE2 has an extra appendage at the N terminal start with the amino acid composition of (1)LDPGLQPGQFSAD(13). Where, LEU1, LEU5, SER11 interacted with the GLU63, TRP68 and ASP13. This interaction blocked the RBD domain of ACE1 as compared to ACE2. Quantitative values like solvated area, degree, betweenness centrality and closeness centrality of LEU1, LEU5, SER11 interaction has also been presented in Fig.5, which indicated that, structural involvement of this extra appendage with RBD domain. The RBD domain of ACE2 remain uncovered as according to the PDB structures 1r4l, 1r42, 3d0h, 3d0i, 3scj, 3sck, 3scl, 7kj3 and 7kj4. Whereas structure of ACE1 (6en5) showed the four unit with that extra appendage. So, nCOV2 spike attachment to ACE2 has been facilitated due to the hassle less interaction in comparison to ACE1.

**Figure 5.**
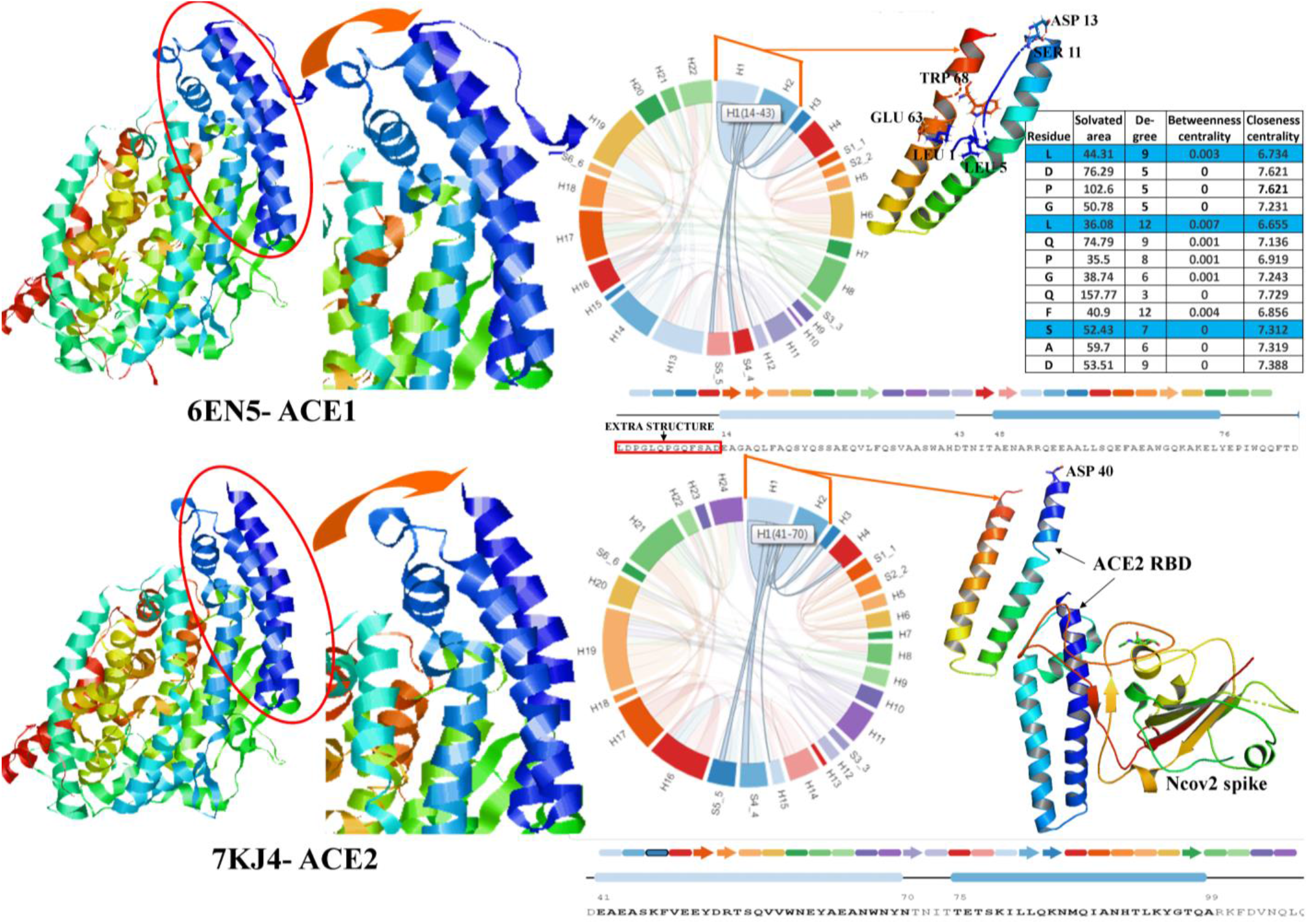
Comparative study between ACE1 and ACE2 receptor binding domain (RBD). Though they have sequence, secondary and tertiary structure level similarity, ACE1 have extra appendage which interacted with the RBD domain with amino acids combination of LEU1-GLU63, LEU5-TRP68 and SER11-ASP13, which blocks the nCOV spike attachment with ACE1. Whereas, ACE2 has no extra part which facilitated the nCOV2 spike binding.

According to Ali and Vijayan (2020) [17], mechanism of nCOV2-ACE2 interaction starts near the N-terminus of ACE2 with the amino acids ofAsp38, Glu35 and Lys353 which has also decided in Figure1. So, in our study, selection of CUT segment throughout the RBD domain is significant rather than a partial one. This will confirm the complete detachment of nCOV spike binding. Report revealed that synthetic inhibitor comprising two α helical peptides homologues to protease domain of ACE2 can block the RBD domain of S protein of the SARS CoV 2. This significant docking may have some therapeutic implications also. Similar type of study suggests that engineered ACE2 receptor traps can neutralize SARS-CoV-2[18]. A hexapeptide in core of spike RBD inhibits the association between spike S1 and ACE-2 [19].

This work has been shown in cultured human cells and in experimental animal model. But as this peptide is a very short fragment having no higher-order of protein structure so the effect may not be due to the complementarily based S1 and ACE-2 binding. Several organ-protecting effects of this hexapeptide justify more intense work in this aspect [19]. However, in the current study we demonstrated that in-silico specific amino acid alteration in the highly conserved S protein of the SARS CoV 2 can potentially block its ACE2 binding, hence minimize the viral infection. The present study has immense therapeutic implication in Covid-19 research. The spike proteins of the SARS CoV 2 from the 186 countries of origin were tested and the different conserved fraction and conserved amino acids were targeted for mutation to test the ACE2 blocking effects. It was noticed that T500S, Y489S,T500S and Y489S,Y453S,T500Y could be a great universal ACE2 blocker, this work has highly promising therapeutic applications. Further studies are necessary.

